# Disentangling the representation of identity from head view along the human face processing pathway

**DOI:** 10.1101/045823

**Authors:** J. Swaroop Guntupalli, Kelsey G. Wheeler, M. Ida Gobbini

## Abstract

Neural models of a distributed system for face perception implicate a network of regions in the ventral visual stream for recognition of identity. Here, we report an fMRI neural decoding study in humans that shows that this pathway culminates in a right inferior frontal cortex face area (rIFFA) with a representation of individual identities that has been disentangled from variable visual features in different images of the same person. At earlier stages in the pathway, processing begins in early visual cortex and the occipital face area (OFA) with representations of head view that are invariant across identities, and proceeds to an intermediate level of representation in the fusiform face area (FFA) in which identity is emerging but still entangled with head view. Three-dimensional, view-invariant representation of identities in the rIFFA may be the critical link to the extended system for face perception, affording activation of person knowledge and emotional responses to familiar faces.

**Significance Statement:** In this fMRI decoding experiment, we address how face images are processed in successive stages to disentangle the view-invariant representation of identity from variable visual features. Representations in early visual cortex and the occipital face area distinguish head views, invariant across identities. An intermediate level of representation in the fusiform face area distinguishes identities but still is entangled with head view. The face-processing pathway culminates in the right inferior frontal area with representation of view-independent identity. This paper clarifies the homologies between the human and macaque face processing systems. The findings show further, however, the importance of the inferior frontal cortex in decoding face identity, a result that has not yet been reported in the monkey literature.

## Introduction

Humans arguably can recognize an unlimited number of face identities but the neural mechanisms underlying this remarkable capability are still unclear. The human neural system for face perception (1–4) consists of distributed cortical fields for the visual analysis of faces, and the computations performed in this system are a matter of intense investigation and controversy. Freiwald and Tsao (5) analyzed the neural population responses in cortical face patches in macaque temporal lobes identified with fMRI.

While population codes in the more posterior face areas, ML and MF, represent face view that is invariant across identities, population codes in the most anterior face-responsive area AM represent face identity that is almost fully view-invariant. A face patch located intermediately (AL) is tuned to mirror symmetric views of faces.

Here we show, for the first time, a progressive disentangling of the representation of face identity from the representation of head view in the human face processing system with a structure that parallels that of the macaque face patch system (5). While early visual cortex (EVC) and the occipital face area (OFA) distinguish head views, view-invariant representation of identities in the human face perception system is fully achieved in a right inferior frontal face area (rIFFA). The representation of faces in the fusiform face area (FFA) revealed an intermediate stage of processing at which identity begins to emerge but is still entangled with head view.

## Results

Subjects viewed four previously unfamiliar identities, two male and two female, with five different head views: left and right full profiles, left and right half profiles, and frontal view. Subjects were visually familiarized with the identities a day before scanning by watching videos of each identity and then performing an identity matching task to facilitate further visual learning. In each trial, the same face image was presented three times in rapid succession with small variations in its size and location (Fig. 1). Subjects performed a one-back identity repetition detection task to ensure attention to the stimuli. We performed multivariate pattern classification and representational similarity analyses across the whole brain using searchlights and in face-selective regions of interest (ROIs).

**Fig. 1.**
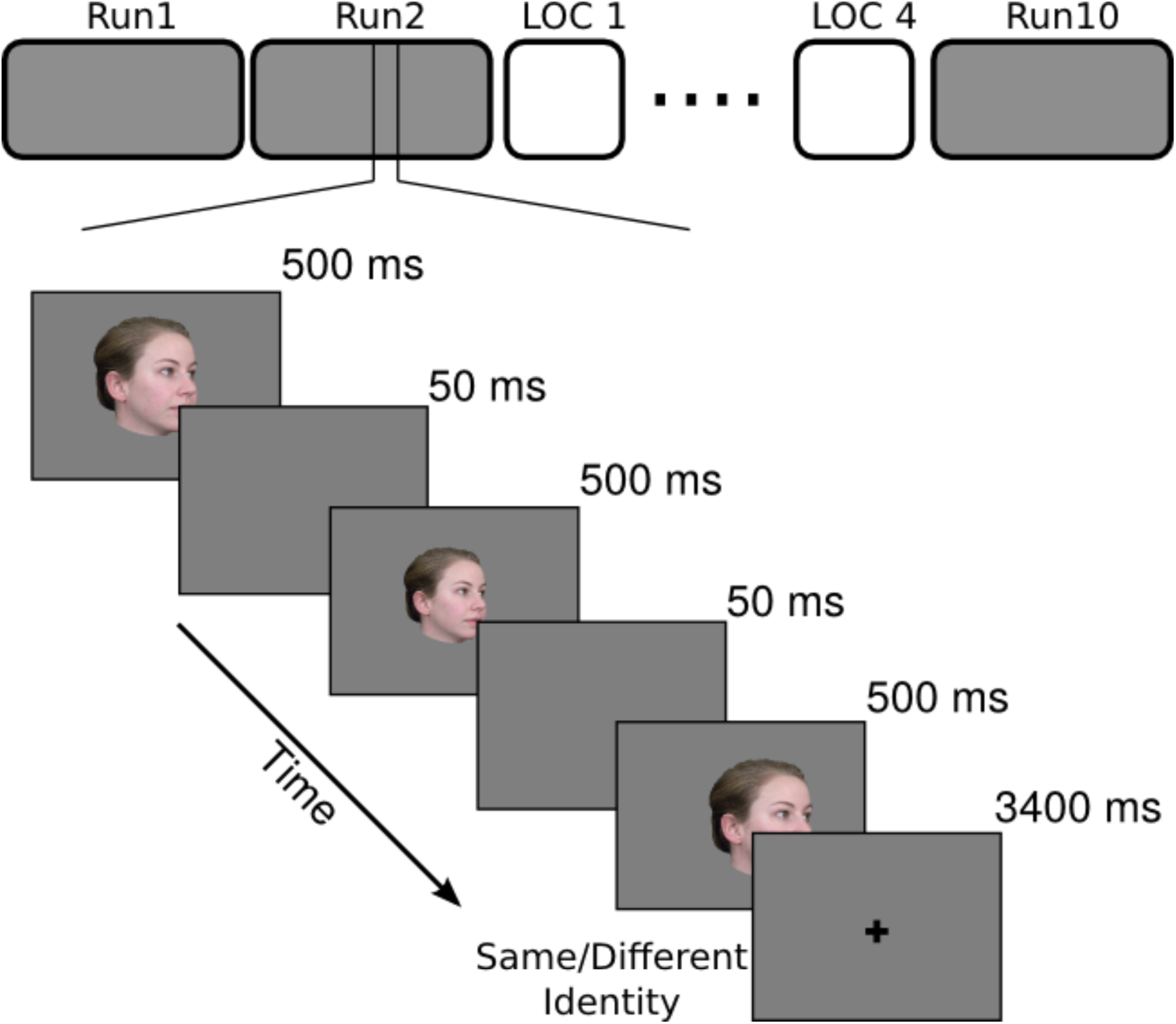
Schematic of the main experiment with an example of the trial structure. fMRI study had 10 runs of the main experiment and 4 runs of localizer interspersed. Each run of the main experiment had 63 trials - 60 stimulus trials and 3 fixation trials. Each trial started with one of the 20 face images presented three times at variable size and location with 50 ms of ISI between presentations, and ended with a 3400 ms fixation. Subjects performed a 1-back identity matching task to keep their attention to the stimuli.

### Classification analyses

First, we used multivariate pattern classification (MVPC) in surface-based searchlights to identify cortical areas that encode faces in terms of view and identity (6). For classification of identity invariant over face views, a classifier was trained to classify four identities over 4 views and tested on the left-out view of the four identities. For classification of head view invariant over identities, a classifier was trained to classify head views over three identities and tested on five head views of the left-out identity. Results (Fig. 2A) show robust representation of face identity invariant to head view in the right inferior frontal cluster, the rIFFA, with peak classification accuracy of 41.2% (chance = 25%), and representation of view invariant to face identity in a large expanse of early visual cortex (EVC) that includes the occipital face area (OFA) and part of the right fusiform face area (FFA) with peak classification accuracy of 65.4% (chance = 20%). To visualize the representational geometries of responses to face images with different identities and head views, we performed a multi-dimensional scaling analyses (MDS) of patterns response in these two clusters. MDS of the head view cluster clearly shows a representation of faces arranged according to head view invariant to identity with a circular geometry in which adjacent head views are closer to each other and the left and right full profiles are closer to each other (Fig. 2B). In contrast, MDS of the rIFFA identity cluster shows a representation of faces arranged according to identity invariant to view with the responses to each of the four identities clustered together for most or all head views.

**Fig. 2.**
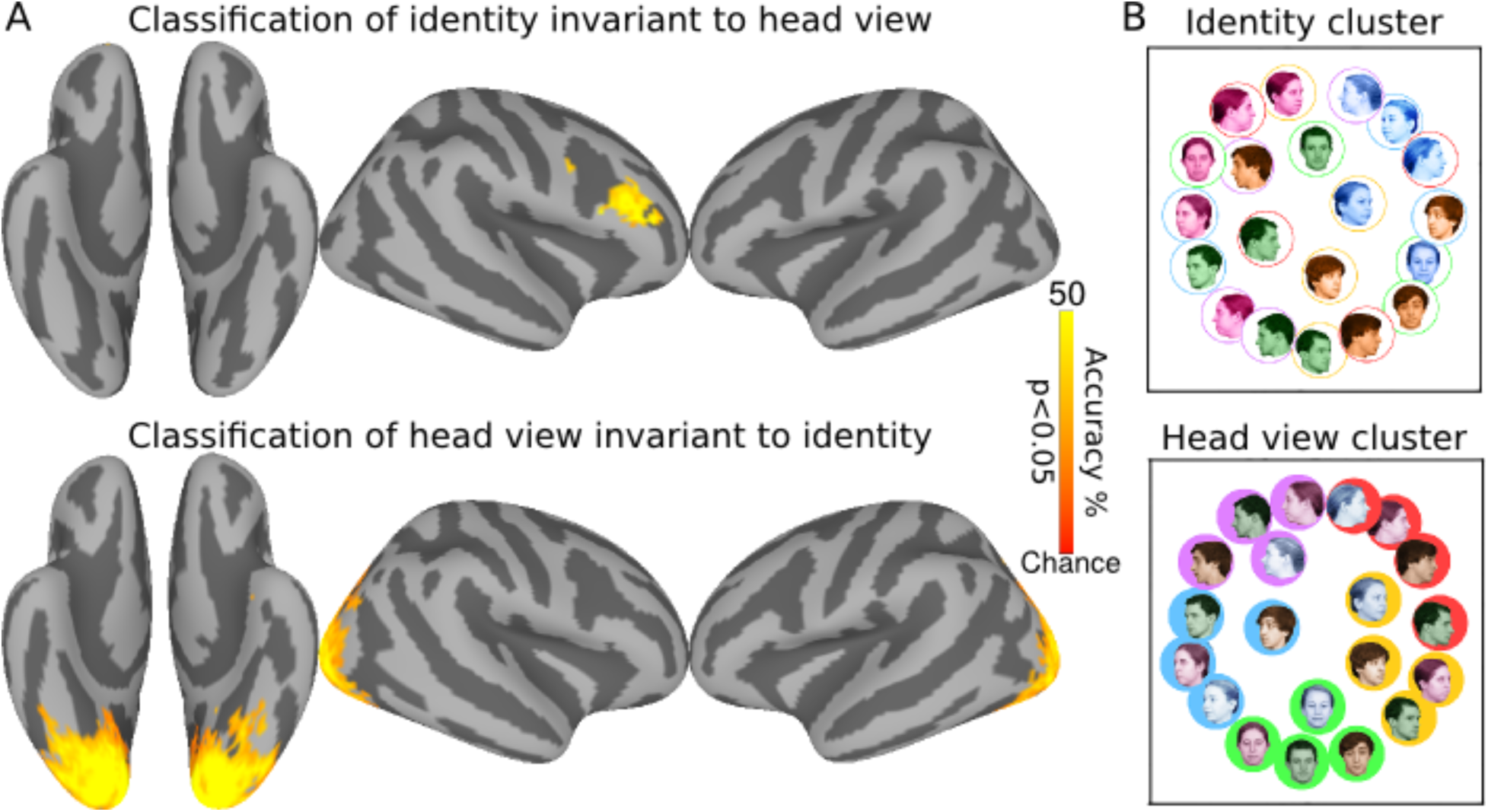
Surface searchlight classification of faces. (A) Classification accuracies for face identity cross-validated over views (*top*) and head view cross-validated over identities (*bottom*). Chance accuracy is 25% for face identity and 20% for head view classifications. Maps are thresholded at p<0.05 after correcting for multiple comparisons using permutation testing. (B) MDS plots of representational geometries of responses to face stimuli in the identity cluster (*top*) and the head view cluster (*bottom*).

### Representational similarity analyses

For an analysis of representational geometry across cortex, we next performed a searchlight representational similarity analysis (RSA) (7). We constructed three model similarity structures capturing representation of 1) identity invariant to view, 2) mirror symmetry of views, and 3) view invariant to identity (Supplementary Fig. 1). We performed ridge regression to fit these three model similarity structures in surface-based searchlights, producing three coefficients in each searchlight. Fig. 3 shows cortical clusters with coefficients for models capturing identity invariant to views and views invariant to identity. Consistent with the classification results, representational geometry in the right inferior frontal cluster is correlated significantly with the face identity similarity model and representational geometry in a large EVC cluster that included OFA and part of the right and left FFA correlated significantly with the head view similarity model.

**Fig. 3.**
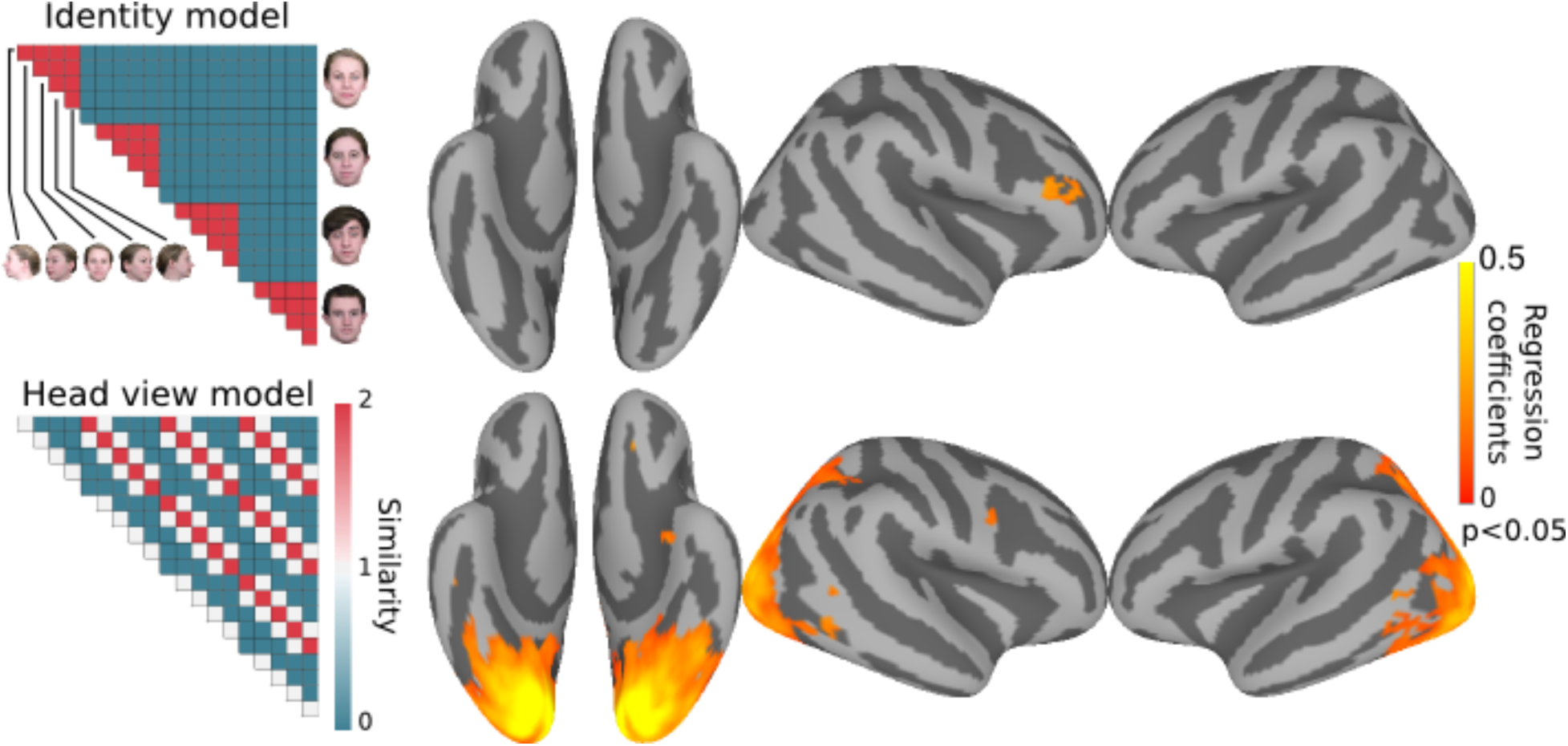
Surface searchlight based modeling of representational geometry. Neural representation of faces in each searchlight was modeled with three model similarity structures as regressors using ridge regression. Representational geometry in the rIFFA correlated with the identity model (*top*), whereas representational geometry in EVC correlated with the head view model (*bottom).* Maps are thresholded at p<0.05 after correcting for multiple comparisons using permutation testing. Correlation with the mirror symmetry model did not reveal any significant clusters.

### Classification analyses and representational similarity analyses in face-selective ROIs

Searchlight analyses revealed representation of head view in posterior visual cortex and identity in the rIFFA but unlike other reports these searchlight analyses did not find representation of identity in ventral temporal (VT) face areas (8–13). To investigate further the representation of faces in the core face system, we defined face-selective regions in all subjects with a localizer. We defined the FFA, OFA, posterior superior temporal sulcus face area (pSTS), and ATFA (Fig. 4A). We then performed MVPC and RSA in each of these ROIs. Classification of face identity, invariant to views, was significant in FFA and ATFA with average accuracies of 36.5% (Chance=25%; 95% CI=[28.5%, 45.8%]) and 30.8% (95% CI=[27.3%, 34.2%]) respectively, but was not different from chance in OFA and pSTS (Fig. 4B). Classification of head view, invariant to identity, was successful in OFA and FFA with average accuracies of 44.2% (Chance=20%; 95% CI=[35.0%, 50.4%]) and 29.2% (95% CI=[20.8%, 38.1%]) respectively, but was not different from chance in ATFA and pSTS (Fig. 4B). Analysis of neural representational geometry showed that representational geometry in the OFA correlated significantly only with head view model (beta=0.33; p<0.001) (Fig. 4C). Representational geometry in the FFA correlated significantly with head view (beta=0.12; p<0.001), mirror symmetry (beta=0.08; p<0.01), and identity models (beta=0.16; p<0.001) corroborating its intermediate role in disentangling identity from head view and mirror symmetry (Fig. 4C). Representational geometry in the ATFA did not significantly correlate with any model and representational geometry in the pSTS correlated only with the head view model (beta=0.07; p<0.01).

**Fig. 4.**
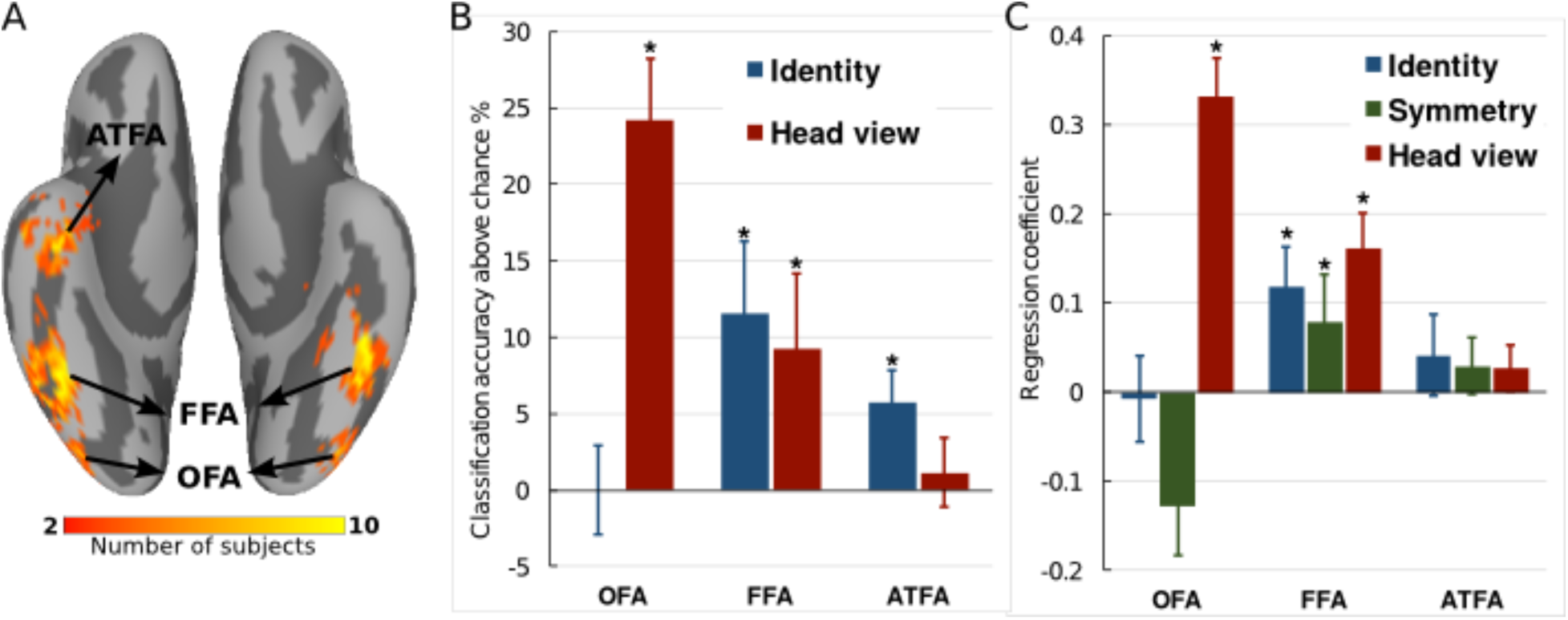
Classification and representational similarity analyses in face-selective ROIs. (A) Anatomical locations of face-selective ROIs as determined by localizer. (B) Classification accuracies of face identity invariant to view and head view invariant to identity in regions of the core face system. (C) Modeling representational geometry in ROIs. Asterisks indicate accuracies that were significant with permutation. Error bars indicate standard error (SEM).

## Discussion

Our results show a progressive disentangling of the representation of face identity from the variable visual features of different images of that face in a hierarchically-organized distributed neural system in occipital, VT, and inferior frontal human cortices. Unlike previous reports (5, 8–12) we show that this disentangling process culminates in a face area in right inferior frontal cortex, the rIFFA. The representational geometry of rIFFA responses to face images grouped images by identity and not by head view. By contrast, the representational geometry in EVC and the OFA grouped the same images by head view and not by identity. At an intermediate stage in the FFA, representational geometry reflected both identity and head view. A view-invariant representation of identity also was found in the ATFA but was not as strong as the representation in the rIFFA. Our results reveal an organization similar to that described in monkeys based on single unit recording (5) but these studies did not examine population codes in the frontal face patch.

A face-responsive area in the inferior frontal cortex was first reported in humans using functional brain imaging and monkeys using single unit recording (13–18). Further reports of this area followed in fMRI studies in humans (12, 19–22) and monkeys (20, 23). The human neuroimaging studies have found this area to be face-responsive using perceptual matching of different views of the same identity (13), face working memory (14, 15, 17), retrieval from long-term memory (16), imagery from long-term memory (19), repetition-suppression (24), release from adaptation (26), and functional localizers with dynamic face stimuli (21, 22). The existence of face selective neurons in the inferior frontal cortex also was shown in a human patient with implanted electrodes who reported face-related hallucinations after direct stimulation in prefrontal cortex (27).

Previous fMRI studies of identity decoding using multivariate pattern classification, however, have mostly concentrated on the ventral visual pathway in temporal cortex, using imaging volumes or ROIs that excluded frontal areas (8–11), with the exception of a recent report (12). Previous identity decoding studies found identity information in the posterior VT cortex including the FFA (9, 11, 12) and anterior temporal areas, albeit with locations that are inconsistent across reports (8–11) and absent in one (12). None of these reports analyzed the representation of face view or how the representation of identity is progressively disentangled from the representation of face view in the face processing system. In separate studies, representation of head view and mirror-symmetry have been reported in more posterior locations both in face responsive areas such as the OFA but also in areas that are object-responsive such as the parahippocampal gyrus and in the dorsal visual pathway (9–10, 24, 25). We find that view-invariance of the representation of identity in FFA is limited as it is entangled with the representation of face view, including mirror symmetry, suggesting that it may be more like the monkey face patch AL than ML/MF. Identity-invariant representation of head view in EVC and OFA suggests that ML/MF may be more like the OFA than FFA. The rIFFA appears to be difficult to identify with localizers that use static images without multiple views of the same identity. It is more consistently activated by tasks that involve matching identity across views, dynamic images, or face memory. The increased sensitivity to dynamic face stimuli led Duchaine and Yovel (4) to conclude that this area is part of a “dorsal face pathway” that is more involved in processing face movement, but the review of the literature and our current results suggest that this area plays a key role in the representation of identity that is integrated across face views. Dynamic stimuli may enhance the response in this area because they present changing views of the same identity in a natural sequence. Dynamic visual features that capture how face images change with natural movement may play an important role in building a three-dimensional view-invariant representation (28, 29). Prior to scanning subjects saw dynamic videos of the four identities to afford learning a robust, three-dimensional representation of each identity.

We were able to replicate others’ findings in anterior temporal cortex generally, but only with the ROI analysis. Identity decoding accuracy in the ATFA was lower than accuracy in the FFA, but that may be due to the larger number of voxels in the FFA ROI and the greater reliability of identifying face-selective voxels there. The rIFFA was identified initially with multivariate pattern analyses but also showed face-selectivity with univariate contrasts (Fig. S2), whereas the ATFA was identified only with the localizer. Both the ATFA and rIFFA showed significant decoding of identity and no trend towards decoding head view. The nature of further processing that is realized in the rIFFA, and how this area interacts with the ATFA, remains unclear. Single unit recording studies of the representation of identity in the monkey inferior frontal face patch may further elucidate the role of this face-patch in representing face identity. Such studies, however, may require familiarization with face identities using dynamic stimuli and/or a task that involves memory.

A view-invariant representation of a face’s identity may be necessary to activate person knowledge about that individual and evoke an appropriate emotional response (2, 29).

Thus, the view-invariant representation in rIFFA may provide a link to the extended system for face perception, most notably regions in medial prefrontal cortex and temporoparietal junction for person knowledge and the anterior insula and amygdala for emotion (1–3), and thereby be critical for engaging the extended system in the successful recognition of familiar individuals.

## Materials and Methods

### Subjects

We scanned 13 healthy right-handed subjects (6 females; mean age = 25.3 ± 3.0) with normal or corrected-to-normal vision. Participants gave written informed consent and the protocol of the study was approved by the local ethical committee.

### Stimuli

Four undergraduates (2 females) from Dartmouth College served as models for face stimuli. We took still pictures and short videos of each model. Silent videos were of the head and shoulders, included natural movements to the right and left, as well as muted interactions with the experimenter, and were 15 seconds each. Still face images were color pictures taken with 5 different head views: left and right full profile, left and right half profile, and full-frontal view. To assure consistent image quality, all pictures were made in the same studio with identical equipment and lighting conditions. All still images were cropped to include the hair. Each image was scaled to resolution of 500x500 pixels. Before training, we confirmed that participants did not know any of the identities shown in the experiment.

### Training Session before the fMRI experiment

Subjects were visually familiarized with the identities of the four stimulus models on the day before the scanning session through a short training held in our laboratory. Subjects passively watched a 15s videos without audio of each identity then performed a face identity matching task in which they saw two stimuli in succession with 0.5 s interstimulus interval. Stimuli were still images of the four models at different head views (presented for 1 s) or 2 s video clips. Subjects indicated if the identity was the same or different using a keyboard. There were 240 trials in total, with matching identity in half.

### Scanning

During scanning, each subject participated in 10 functional runs of the main experiment. Each run had 63 trials (60 stimulus trials and 3 fixation trials), and started and ended with a 15s of fixation of black cross on gray background. Each stimulus trial was 5 s long and started with a stimulus image presented for 500 ms followed by a 50 ms gray screen, repeated three times, followed by 3400 ms of fixation. The three repetitions on each trial were of the same identity and head view but with the image size and location jittered (+/-50 pixels equivalent to 1.25 degrees variations in image size and +/-10 pixel variations in the horizontal and vertical location). Each face image subtended approximately 12.5 degrees of visual angle. Subjects performed a one-back repetition detection task based on identity, pressing a button with the right index finger for ‘same’ and the right middle finger for ‘different’.

### Localizer

In addition to the main experiment, four runs of a functional localizer were interleaved with the experimental runs. Each localizer run had 2 blocks each of faces, objects, and scenes with 8 s of fixation separating them. During each presentation block, subjects saw 16 still images from a category with 900 ms of image presentation and 100 ms of ISI. Subjects performed a one-back repetition detection task. Each run started with 12 s of fixation at the beginning and ended with 12 s of fixation. None of the faces used for the localizer were part of the set of stimuli used for the fMRI experiment on head view and identity.

### fMRI protocol

Subjects were scanned in a Philips Intera Achieva 3T scanner with an 32 channel head coil at the Dartmouth Brain Imaging Center. Functional scans were acquired with an echo planar imaging sequence (TR=2.5 s, TE=30 ms, flip angle=90°, 112 × 112 matrix, FOV=224 mm × 224 mm, R-L phase encoding direction) every 2.5 s with a resolution of 2 ☓ 2 mm covering the whole brain (49×2 mm thick interleaved axial slices). We acquired 140 functional scans in each of the 10 runs. We acquired a fieldmap scan after the last functional run and a T1-weighted anatomical (TR=8.265 ms, TE=3.8 ms, 256 × 256 ☓ 220 matrix) scan at the end. The voxel resolution of anatomical scan was 0.938 mm × 0.938 mm × 1.0 mm.

### Data preprocessing

Each subject’s fMRI data was preprocessed using AFNI software (31). Functional data was first corrected for the order of slice acquisition and then for the head movement by aligning to the fieldmap scan. Functional volumes were corrected for distortion using fieldmap with FSL-Fugue (32). Temporal spikes in the data were removed using 3dDespike in AFNI. Time series in each voxel was filtered using a high-pass filter with a cutoff at 0.00667 hz, and the motion parameters were regressed out using 3dBandpass in AFNI. Data were then spatially smoothed using a 4 mm full-width at half-max Gaussian filter (3dmerge in AFNI). We ran a GLM analysis to estimate beta and t-statistic values for each of the 20 stimulus images using TENT function of 3dDeconvolve in AFNI resulting in 7 estimates from 2.5 s to 17.5 s after stimulus onset. In the end, we obtained 7 estimates per voxel per stimulus image in each subject. We used the first 5 of those 7 response t-statistic estimates in all our analyses (33). Thus, in each searchlight or ROI we measured a pattern of response to each stimulus image as a vector whose features were five timepoints for each voxel. We extracted cortical surfaces from the anatomical scans of subjects using FreeSurfer (34), aligned them to the FreeSurfer’s cortical template, and resampled surfaces into a regular grid with 20,484 nodes using MapIcosahedron in AFNI. We implemented our methods and ran our analyses in PyMVPA (35) unless otherwise specified (http://www.pymvpa.org).

### Definition of face-selective ROIs

We used the same preprocessing steps for face localizer data and estimated the contrast for faces greater than objects using GLM analysis to define face-selective regions. Clusters of voxels with stronger responses to faces were assigned to OFA, FFA, pSTS, and ATFA regions based on their anatomical locations (1). We used faces greater than objects contrast with a threshold of t=2.5 to 3 for bilateral fusiform (FFA), occipital (OFA) and right pSTS face areas, and 2-2.5 for right anterior temporal face area (ATFA). For ATFA and pSTS face area, only the right hemisphere yielded robust ROIs in all subjects, whereas FFA and OFA were identified bilaterally in all subjects but one who had no identifiable OFA at the used threshold range. These criteria based on responses to still images of unfamiliar faces did not identify a consistent face-selective cluster in the rIFFA even though MVPA (classification and similarity analyses) did reveal such a cluster based on face-identity-selective patterns of response. Post-hoc analysis of responses to the localizer stimuli in a searchlight centered on the cortical node with peak accuracy for classification of identity confirmed that also the rIFFA showed face-selectivity based on the univariate contrast between faces and objects (Fig. S2). Table S1 lists the average number of voxels and volumes of face-selective ROIs.

## Multivariate pattern classification (MVPC)

### Searchlight (MVPC)

We performed MVPC analyses to localize representations of face stimuli in terms of head view, independent of identity and identity independent of head view, using surface-based searchlights (36). We centered searchlights on each surface node, and included all voxels within a cortical disc of radius 10 mm. Thickness of each disc was extended by 50% into and outside the gray matter to account for differences between EPI and anatomical scans due to distortion. Identity classifications were performed using a leave-one-viewpoint-out cross validation scheme, and viewpoint classifications were performed using a leave-one-identity-out cross validation scheme. For example, each classifier for identity was built on responses to all head views but one (16 vectors −4 head views of 4 identities) and tested on the left out head view (1 out of 4 classification). This was done for all 5 data folds on head view. Similarly, each classifier for head view was built on all identities but one (15 vectors −5 head views of 3 identities) and tested on the left out identity (1 out of 5 classification). MVPC used a linear support vector machine (SVM) classifier (37). The SVM classifier used the default soft margin option in PyMVPA that automatically scales the C parameter according to the norm of the data. Classification accuracies from each searchlight were placed into their center surface nodes resulting in one accuracy map per subject per classification type. We also performed an identical analysis but using permuted labels, 20 per subject per classification type, for significance testing. To compute significant clusters across subjects, we performed a between-subject threshold-free cluster enhancement procedure (38) using our permuted label accuracy maps. We then thresholded the average accuracy map with correct labels across subjects at t>1.96 (p < 0.05, two-tailed, corrected for multiple comparisons) for visualization.

### Face-selective ROIs

In each face-selective ROI of each subject, we performed classification of identity and head view as described above. We used a nested cross-validation scheme to perform feature selection using ANOVA scores. In each fold, training data is used to compute ANOVA scores and classification using different numbers of top features within the training data. The set of features that gave the best accuracy on training data is then used to classify the test data. Both ANOVA scores and which features to use are computed only from the training dataset. We performed significance testing in each ROI for each classification type using permutation testing using 100 permutations in each subject and sampling them with replacement for 10,000 permutations across subjects to compute the null distribution.

### Representation similarity analyses

We performed a representational similarity analysis (7) to model the representational geometry - representational similarity matrices (RSM) - in each searchlight using three models of representation: 1) identity invariant to head view, 2) head view invariant to identity, and 3) head view with mirror symmetry (Fig. S1). Correlation is used to compute similarities between patterns of response to different images. Both the neural and the model similarity matrices are rank ordered before performing a ridge regression with alpha=0.1 to fit the three model similarity structures to the neural similarity structure. Corresponding betas for each model were then collected into the center node for that searchlight. We performed permutation testing to assess the significance of the beta values for each model regression at each surface node using permuted labels and threshold free cluster enhancement method as described above. We then thresholded the average beta maps with correct labels for each model at t>1.96 (corrected) for visualization. We performed a similar modeling of representational geometry in each face-selective ROI. To remove any possible confounds between mirror symmetry and other models, we zeroed out any elements in the mirror symmetry model that overlap with the other two models. We assess the significance of model coefficients in each ROI using permutation testing. We computed 100 model coefficients in each subject using permuted labels, and sampling them with replacement for 10,000 samples across subjects to compute null distribution for each model coefficient in each ROI.

## Multidimensional Scaling

We performed multidimensional scaling in significant head view and identity classification clusters and face-selective ROIs to visualize the representation of face stimuli in each of those regions. Since classification clusters were defined on surface, we aggregated the data from all the voxels that participated in searchlights with their center nodes in those clusters in each subject. Data in each cluster of searchlights and each ROI were reduced to a 20 principal component space (all components) before computing distance matrices to account for variable sizes. Pairwise correlation distance matrices were computed for all 20 face stimuli for each cluster and ROI in each subject. Distance matrices were first normalized in each subject by dividing by the maximum correlation distance within that subject and were averaged across subjects to produce an average distance matrix in each cluster and ROI. A metric MDS was performed with 10000 iterations to project the stimuli onto a 2-dimensional space. MDS solutions for face-selective ROIs were also computed using the same procedure. Supplementary Fig. 3 shows MDS plots for these ROIs.

## Acknowledgements

We are grateful to Jim Haxby for helpful discussion and comments on a previous draft of this paper and to Yu-Chien Wu for help with the scanning parameters.

## Supplementary Information

**Fig. S1.**
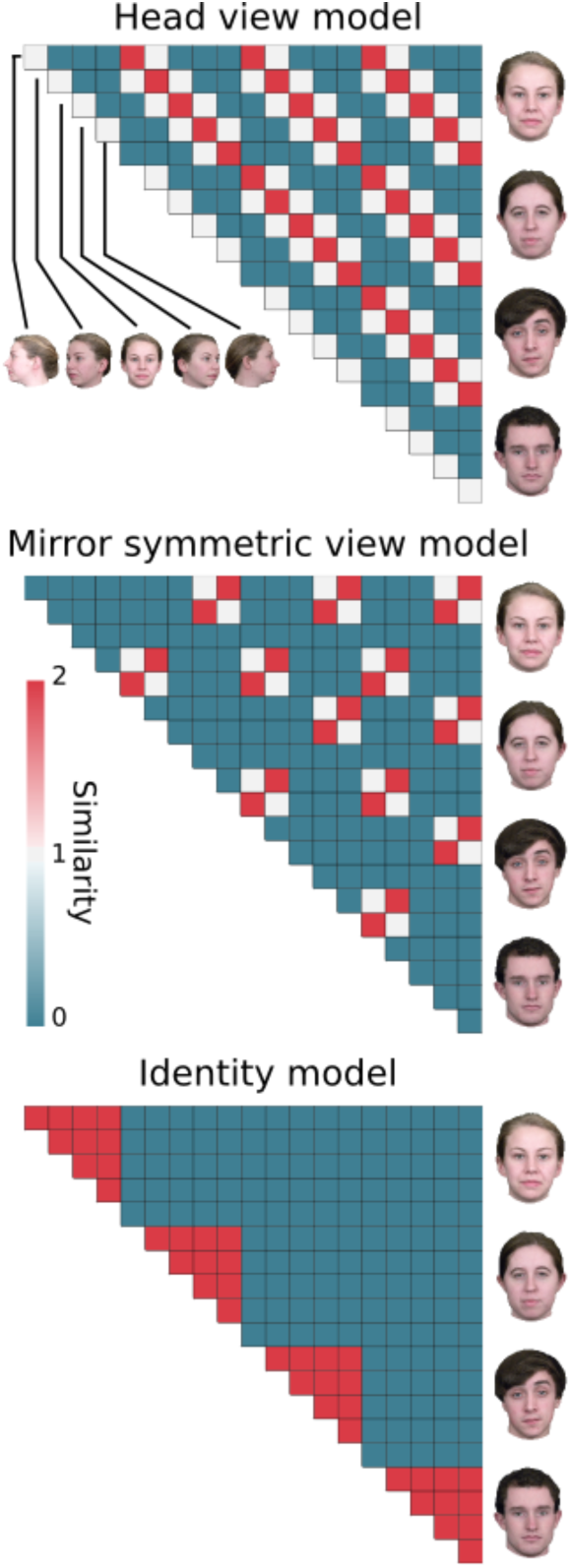
Model similarity structures of face stimuli. Head view model captures similarity between faces based on their view. Mirror symmetric view model captures similarity of face views that are mirror symmetric. Identity model captures similarity of faces based on identity invariant to view.

**Fig. S2.**
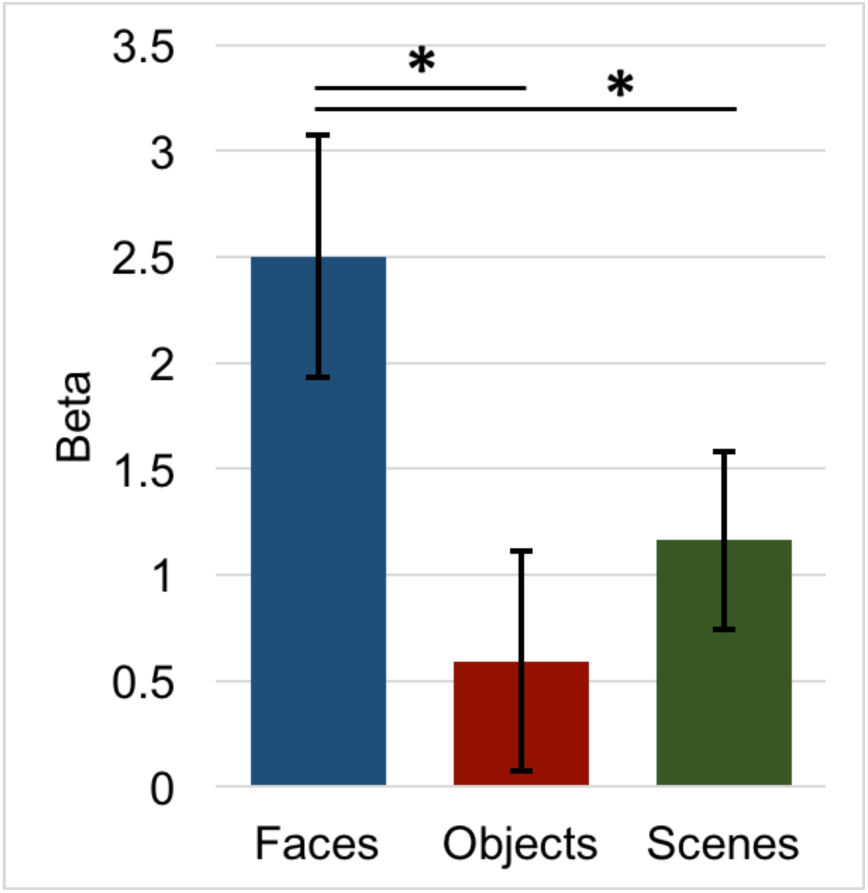
Face selectivity in the right inferior frontal custer. We defined a cortical disc of 10 mm radius centered on the surface node with peak accuracy in the identity classification analysis as our ROI. We then averaged the beta coefficients for each category presented during the localizer in each subject within that ROI. Average estimated response to faces was greater than the response to both objects and scenes across subjects. Error bars indicate SEM, and asterisks indicate p<0.05.

**Fig. S3.**
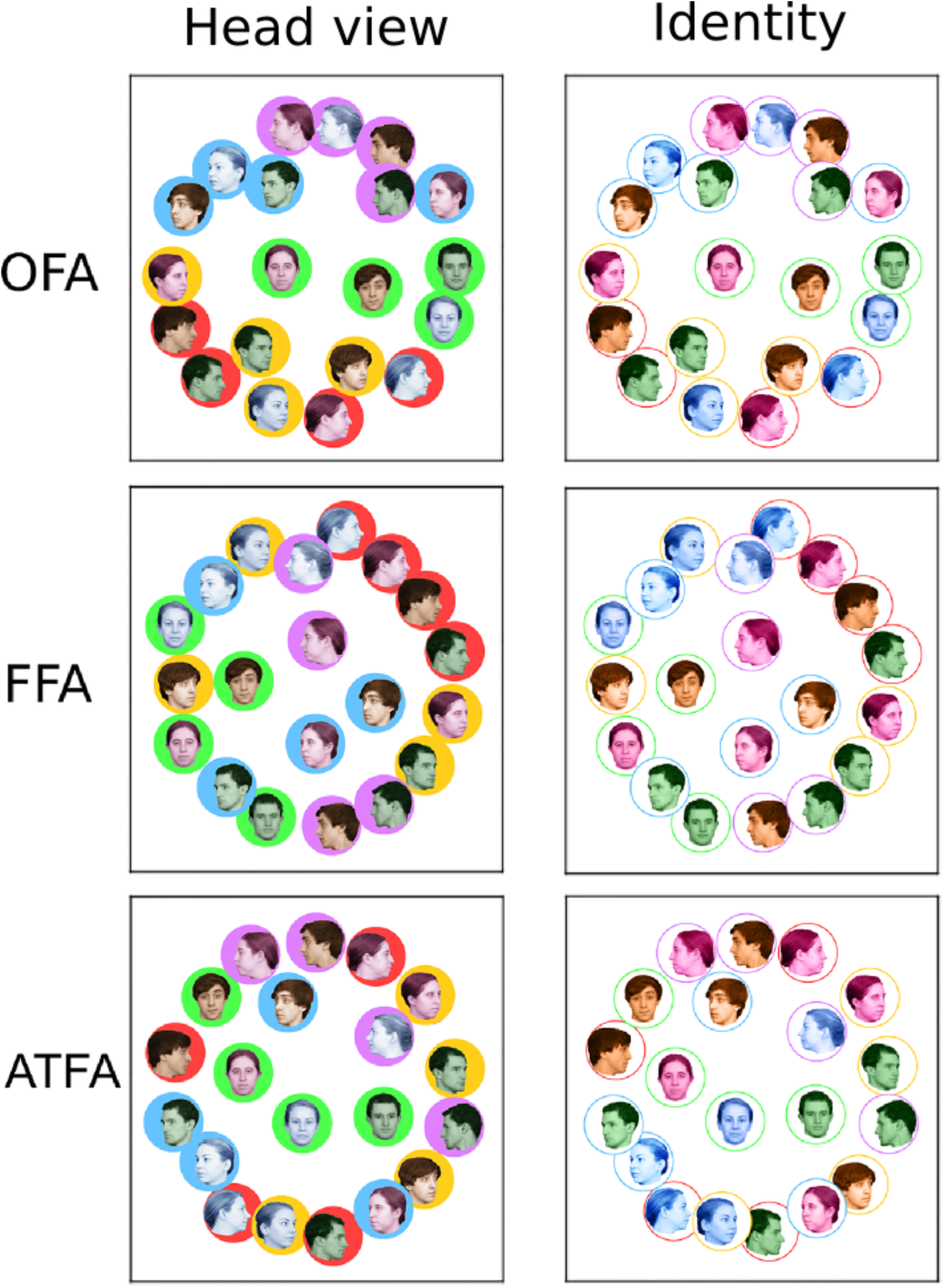
Multidimensional scaling plots of stimulus categories based on the cortical responses in face-selective ROIs. For each ROI, left and right columns depict the same MDS solution with coloring based on head view emphasized on the left and with coloring based on face identity emphasized on the right.

**Table S1.** Size of face-selective regions across subjects

|  | **OFA (RH+LH)** | **FFA (RH+LH)** | **pSTS (RH)** | **ATFA (RH)** |
| --- | --- | --- | --- | --- |
| Voxels | 255.7 | 351.4 | 162.4 | 128.8 |
| Volume (mm<sup>3</sup>) | 2045.3 | 2811.1 | 1299.1 | 1030.8 |

